## Supporting Information for "Delayed postglacial colonization of *Betula* in Iceland and the circum North Atlantic"

### Stóra Viðarvatn Tephrochronology

Each tephra layer was sampled along the vertical axis, sieved to isolate glass fragments between 125 and 500 µm, and embedded in epoxy plugs. Individual glass shards were analyzed at the University of Iceland on a JEOL JXA-8230 election microprobe using an acceleration voltage of 15 kV, beam current of 10 nA and beam diameter of 10 µm. The international A99 standard was used to monitor for instrumental drift and maintain consistency between measurements. Tephra origin was then assessed following the systematic procedures outlined in Jennings et al. (2014) and Harning et al. (2018). Briefly, based on $SiO_2$ wt% vs total alkali ($Na_2O$+$K_2O$) wt%, we determine whether the tephra volcanic source is mafic (tholeiitic or alkalic), intermediate and/or rhyolitic. From here, we objectively discriminate the source volcanic system through a detailed series of bi-elemental plots produced from available compositional data on Icelandic tephra (Harning et al., 2018). Major oxide composition of each tephra layer is provided below in Table S1.

***Table S1:*** *Average and standard deviation major oxide composition (wt%) of Stóra Viðarvatn marker tephra layers used in this study.*

| Depth (cm) | Thickness (cm) | Tephra ID | Age BP | *n* | $SiO_2$ | $TiO_2$ | $Al_2O_3$ | FeO | MnO | MgO | CaO | $Na_2O$ | $K_2O$ | $P_2O_5$ | Total |
|---|---|---|---|---|---|---|---|---|---|---|---|---|---|---|---|
| 342.2 | 0.80 | Hekla 4 | 4200 | 18 | 73.84 | 0.11 | 13.21 | 1.94 | 0.07 | 0.03 | 1.34 | 4.56 | 2.76 | 0.02 | 97.88 |
| | | | | *SD* | *0.79* | *0.03* | *0.20* | *0.10* | *0.04* | *0.05* | *0.08* | *0.16* | *0.08* | *0.02* | *0.93* |
| | | | | 7 | 58.58 | 1.25 | 14.65 | 9.52 | 0.25 | 1.80 | 5.17 | 4.17 | 1.45 | 0.57 | 97.42 |
| | | | | *SD* | *1.21* | *0.17* | *0.26* | *0.38* | *0.04* | *0.37* | *0.29* | *0.19* | *0.09* | *0.12* | *0.59* |
| | | | | 8 | 46.19 | 3.80 | 13.45 | 14.24 | 0.22 | 5.51 | 10.24 | 2.96 | 0.58 | 0.47 | 97.65 |
| | | | | *SD* | *0.30* | *0.47* | *0.75* | *0.38* | *0.05* | *0.44* | *0.44* | *0.11* | *0.10* | *0.26* | *0.40* |
| 414 | 0.50 | Kverkfjöll | 5200 | 8 | 49.98 | 3.10 | 12.98 | 14.71 | 0.23 | 4.43 | 9.02 | 3.02 | 0.67 | 0.39 | 98.52 |
| | | | | *SD* | *0.41* | *0.07* | *0.17* | *0.23* | *0.03* | *0.22* | *0.18* | *0.06* | *0.04* | *0.03* | *0.37* |
| 503.2 | 0.10 | Hekla | 6200 | 10 | 63.61 | 0.74 | 15.15 | 8.57 | 0.24 | 0.90 | 4.15 | 4.03 | 1.78 | 0.29 | 99.44 |
| | | | | *SD* | *1.16* | *0.10* | *0.30* | *0.67* | *0.03* | *0.16* | *0.22* | *0.93* | *0.14* | *0.06* | *0.83* |
| | | Kverkfjöll | | 8 | 49.67 | 2.90 | 13.09 | 14.11 | 0.23 | 5.17 | 9.88 | 2.65 | 0.44 | 0.35 | 98.48 |
| | | | | *SD* | *0.46* | *0.47* | *0.50* | *0.92* | *0.03* | *0.79* | *1.12* | *0.21* | *0.12* | *0.11* | *0.77* |
| | | Askja? | | 2 | 49.86 | 1.75 | 13.97 | 11.81 | 0.18 | 6.71 | 11.88 | 2.27 | 0.21 | 0.17 | 98.79 |
| | | | | *SD* | *0.21* | *0.05* | *0.33* | *0.01* | *0.01* | *0.30* | *0.11* | *0.11* | *0.02* | *0.03* | *0.29* |
| 821 | 31.0 | G10ka Series | 10400-9900 | *353* | 49.51 | 2.82 | 13.30 | 14.05 | 0.23 | 5.68 | 10.10 | 2.67 | 0.42 | 0.31 | 99.09 |
| | | | | *SD* | *0.39* | *0.15* | *0.21* | *0.42* | *0.02* | *0.25* | *0.28* | *0.14* | *0.04* | *0.04* | *0.53* |
| | | | | 2 | 50.66 | 1.82 | 13.80 | 12.92 | 0.25 | 6.25 | 10.94 | 2.54 | 0.32 | 0.18 | 99.66 |
| | | | | *SD* | *0.36* | *0.09* | *0.22* | *0.88* | *0.04* | *0.49* | *0.91* | *0.13* | *0.07* | *0.01* | *0.14* |
| | | | | 1 | 48.99 | 2.65 | 13.01 | 15.78 | 0.24 | 4.93 | 10.20 | 2.74 | 0.34 | 0.22 | 99.09 |
| | | | | 4 | 46.93 | 4.63 | 12.90 | 15.21 | 0.24 | 5.02 | 10.12 | 3.13 | 0.71 | 0.58 | 99.45 |
| | | | | *SD* | *0.51* | *0.64* | *0.54* | *0.79* | *0.06* | *0.41* | *0.59* | *0.10* | *0.12* | *0.09* | *0.34* |
| | | | | 3 | 49.37 | 3.30 | 12.85 | 15.24 | 0.24 | 4.93 | 9.43 | 2.74 | 0.63 | 0.38 | 99.10 |
| | | | | *SD* | *0.64* | *0.08* | *0.21* | *0.25* | *0.01* | *0.47* | *0.67* | *0.16* | *0.14* | *0.07* | *1.19* |
| 885 | 2.85 | Askja S | 10830 | 45 | 74.89 | 0.30 | 12.35 | 2.77 | 0.09 | 0.25 | 1.58 | 4.35 | 2.56 | 0.04 | 99.17 |
| | | | | *SD* | *1.18* | *0.03* | *0.21* | *0.12* | *0.03* | *0.02* | *0.08* | *0.49* | *0.11* | *0.04* | *1.48* |
| | | | | 34 | 50.02 | 1.82 | 13.73 | 12.83 | 0.22 | 6.48 | 11.36 | 2.46 | 0.28 | 0.17 | 99.38 |
| | | | | *SD* | *0.34* | *0.13* | *0.20* | *0.46* | *0.03* | *0.28* | *0.28* | *0.12* | *0.10* | *0.03* | *0.58* |
| | | | | 6 | 48.88 | 2.18 | 13.50 | 13.17 | 0.22 | 6.67 | 11.44 | 2.48 | 0.26 | 0.19 | 98.98 |
| | | | | *SD* | *0.13* | *0.03* | *0.07* | *0.22* | *0.04* | *0.05* | *0.06* | *0.07* | *0.01* | *0.03* | *0.27* |

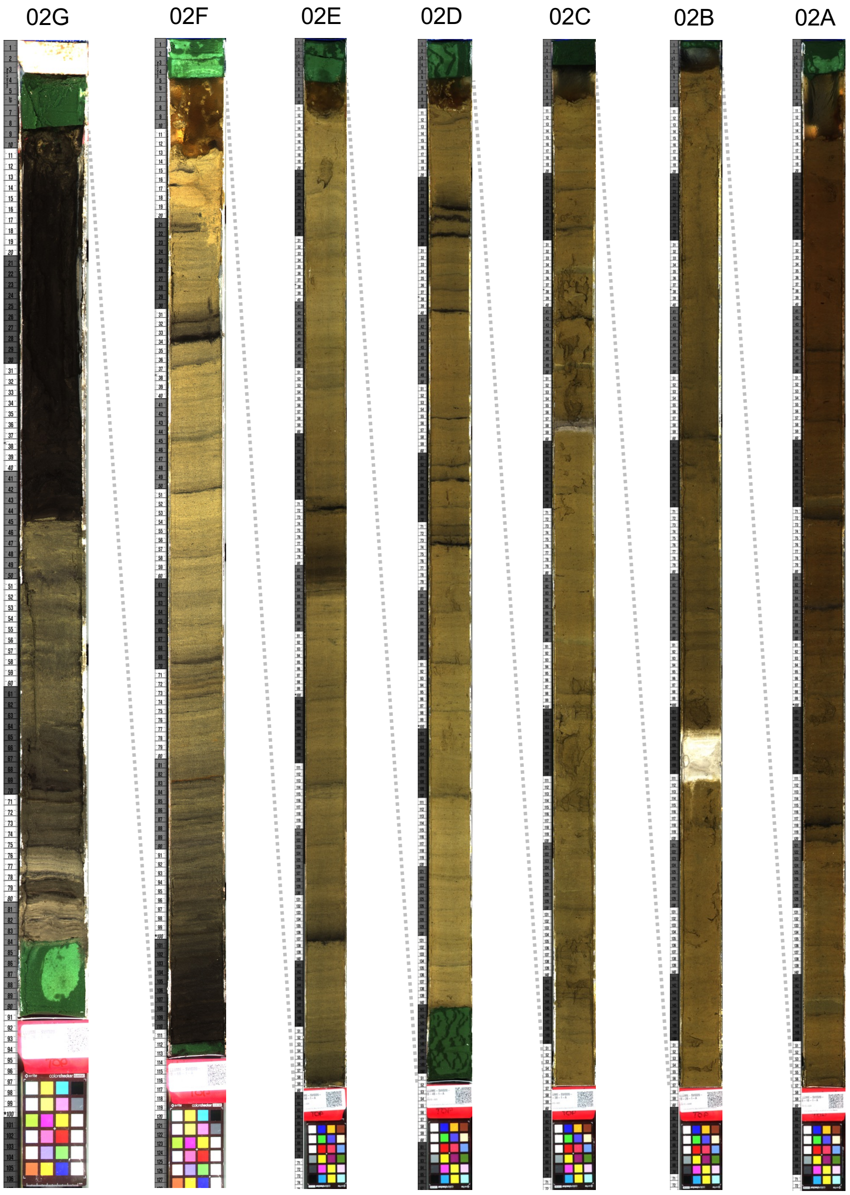


***Figure S1:*** *Core photographs from 20SVID-02 taken at the University of Minnesota's LacCore facility. Section 02G (left) is the bottom and 02A (right) is the top.*

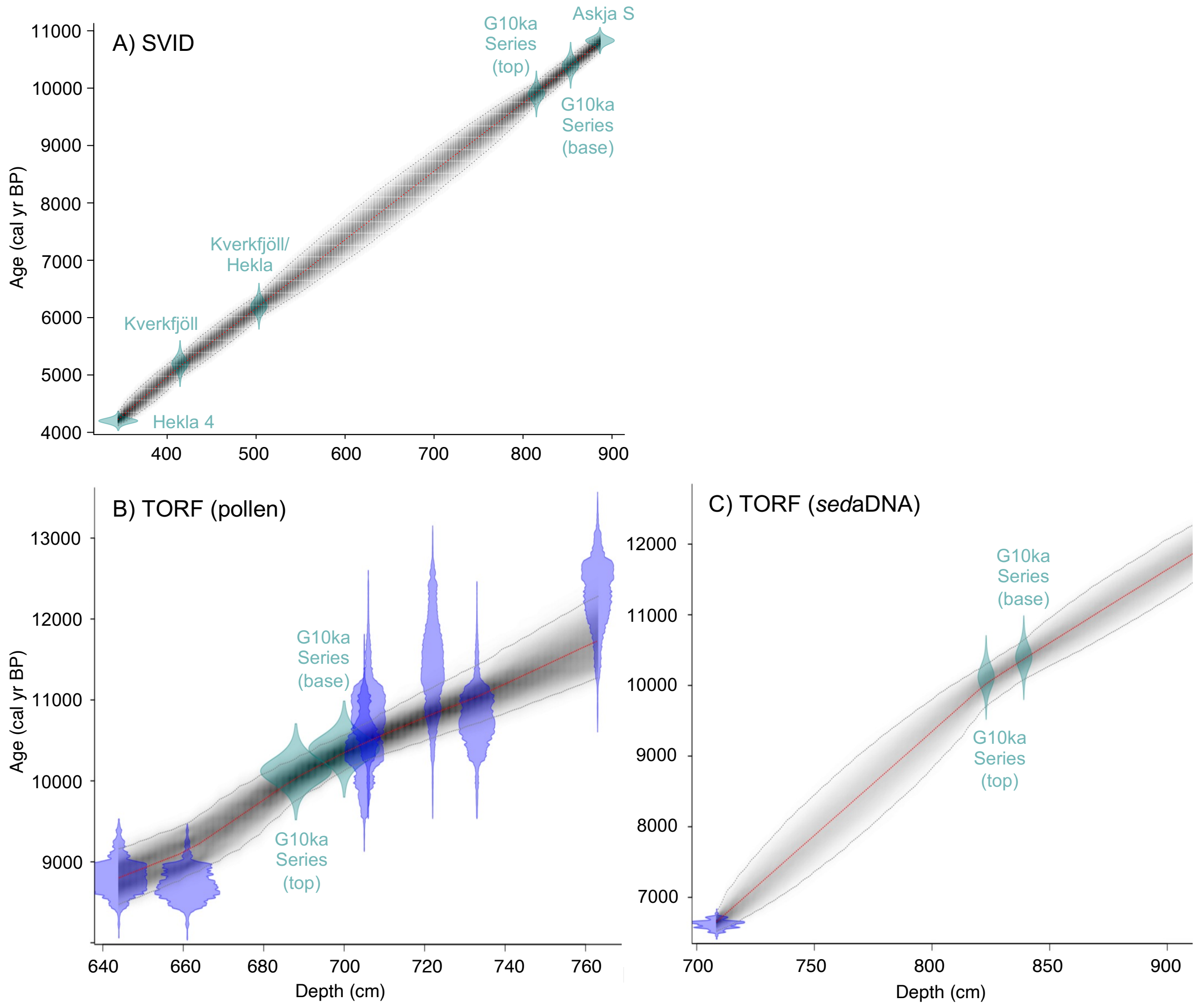


***Figure S2:*** *Iceland lake sediment age models used in this study. A) Stóra Viðarvatn's age model (this study), B) Torfdalsvatn's pollen and macrofossil age model modified from Rundgren (1995, 1998) (see Geirsdóttir et al., 2020), and C) Torfdalsvatn's sedaDNA age model modified from Alsos et al. (2021). Green points reflect tephra layers and blue points reflect calibrated $^{14}C$ ages.*

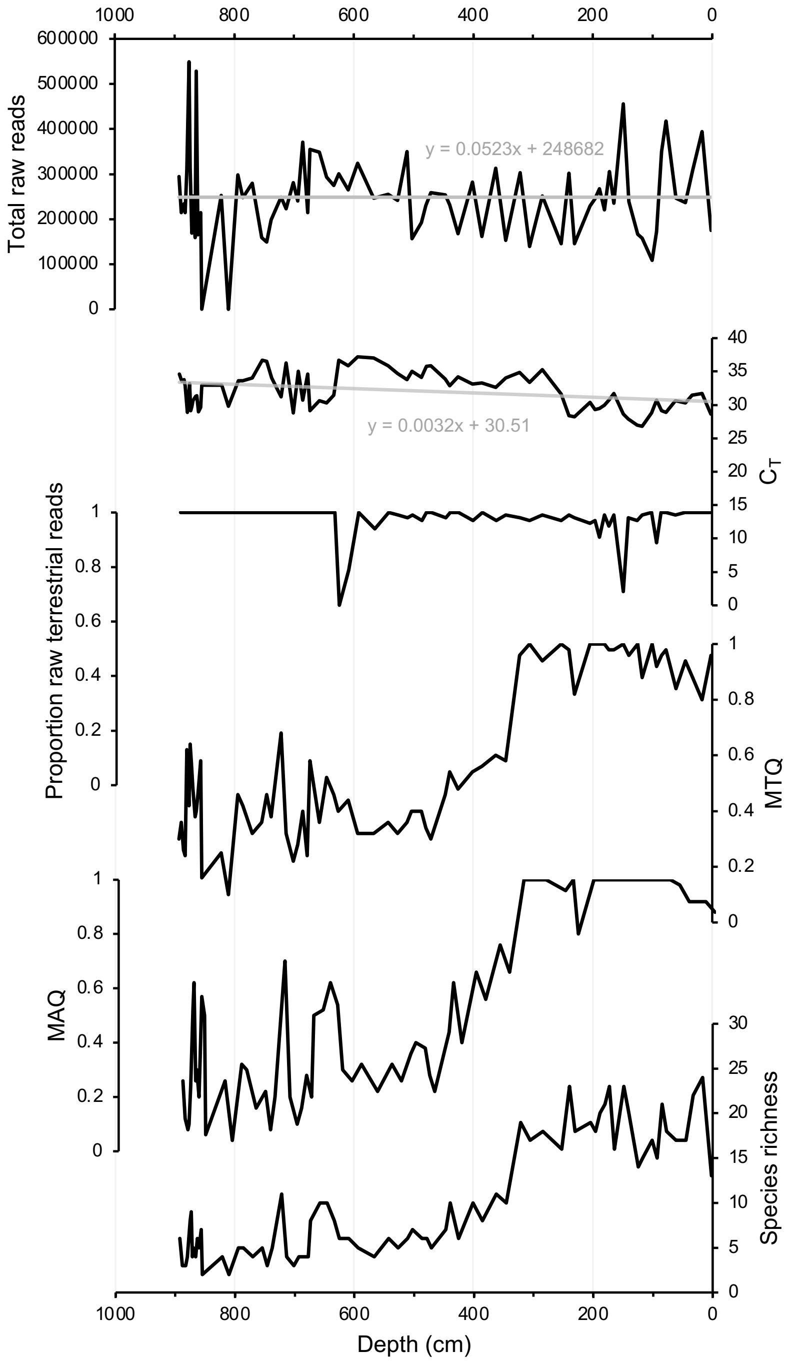


***Figure S3: DNA quality assessment for Stóra Viðarvatn's entire Holocene record plotted against depth (cm).*** *Total raw DNA reads,* $C_T$ *values, proportion raw terrestrial reads, metabarcoding technical quality (MTQ), metabarcoding analytical quality (MAQ), and species richness. Light gray lines are linear regressions, with equations shown.*

***Table S2***: *Primer Sequences used in metabarcoding and qPCR experiments.*

| Primer | Sequence 5’-3’ |
|---|---|
| truseq_trnL_g | ACACTCTTTCCCTACACGACGCTCTTCCGATCTGGGCAATCCTGAGCCAA |
| truseq_trnL_h | GTGACTGGAGTTCAGACGTGTGCTCTTCGATCT TTGAGTCTCTGCACCTATC |

***Table S3:*** *ASV sequences identified in our samples for Salicaceae and Betulaceae.*

| Taxon | ASV Sequence |
|---|---|
| Salicaceae | ATCCTATTTTTCGAAAACAAACAAAGGTTCATAAAGACAGAATAAGAATACAAAAG |
| Betulaceae | ATCCTGTTTTCCGAAAACAAATAAAACAAATTTAAGGGTTCATAAAGTGAGAATAAAAAAG |

***Table S4:*** *Total read data remaining after each data filtering step.*

| Step | Total read data remaining |
|---|---|
| Raw | 18,653,132 |
| Minimum 10 reads per PCR replicate, occurred in 2 of out 5 PCR replicates, and minimum 100 reads across PCR replicates | 17,303,887 |
| Non-native taxa and blank contaminants | 15,909,615 |
